# Expansive and diverse phenotypic landscape of field *Aedes aegypti* larvae with differential susceptibility to temephos: beyond metabolic detoxification

**DOI:** 10.1101/2021.06.07.447310

**Authors:** Jasmine Morgan, J. Enrique Salcedo-Sora, Omar Triana-Chavez, Clare Strode

## Abstract

Arboviruses including dengue, Zika and chikungunya are amongst the most significant public health concerns worldwide and their control relies heavily on the use of insecticides to control the vector mosquito *Aedes aegypti*. The success of controlling these vector-pathogen systems is threatened by widespread insecticide resistance. The work presented here profiled the gene expression of the larvae from two field populations of *Ae. aegypti* with differential susceptibility to temephos. The contrasting phenotypes originated from two Colombian urban locations, Bello and Cúcuta, that we have previously reported to have distinctive disease incidence, socioeconomics, and climate. The closeness of the geographical origin of the study populations was suspected to be highly influential in the profiling of the gene expression of resistance since the mosquito’s resistance levels themselves are highly dependent upon environmental variables. We demonstrated that an exclusive field-to-lab (*Ae. aegypti* reference strain New Orleans) comparison generates an over estimation of differential gene expression (DGE) and that the inclusion of a geographically relevant field control, as used here, yields a more discrete, and likely, more specific set of genes. The composition of the obtained DGE profiles is varied, with commonly reported resistance associated genes such as detoxifying enzymes having only a small representation. We identify cuticle biosynthesis, ion exchange homeostasis, an extensive number of long non-coding RNAs, and chromatin modelling among the specifically and differentially expressed genes in field resistant *Ae. aegypti* larvae. It was also shown that temephos resistant larvae undertake further gene expression responses when temporarily exposed to this insecticide. The results from the sampling triangulation approach undertaken here contributes a discrete DGE profiling with reduced noise that permitted the observation of a greater gene diversity. This deeper gene granularity significantly increases the number of potential targets for the control of insecticide resistant mosquitoes and widens our knowledge base on the complex phenotypic network of the *Ae. aegypti* mosquito responses to insecticides.

**Author Summary:** *Aedes aegypti* mosquitoes are vectors for several significant human viruses including dengue, Zika and chikungunya. The lack of widely available vaccines and specific antiviral treatments for these viruses means that the principal method for reducing disease burden is through controlling the vector mosquitoes. Mosquito control relies heavily on the use of insecticides and successful vector control is threatened by widespread insecticide resistance in *Ae. aegypti.* Here, we examined changes in gene expression that occur in temephos resistant populations of *Ae. aegypti* from two field populations in Colombia. We compare gene expression in resistant larvae from Cúcuta with susceptible larvae from Bello and a susceptible laboratory strain of *Ae. aegypti* (New Orleans). We also compare mosquitoes from Cúcuta with and without temephos exposure. We report several differentially expressed genes beyond those usually reported in resistant mosquitoes. We also demonstrate the over estimation in differential gene expression that can occur when field resistant populations are compared against lab susceptible populations only. The identification of new mechanisms involved in the development of insecticide resistance is crucial to fully understanding how resistance occurs and how best it can be reduced.

## Introduction

Arboviral diseases including dengue, Zika and chikungunya are amongst the most significant public health concerns worldwide. The geographical distribution and prevalence of these arboviruses has been increasing rapidly in recent years with the number of dengue infections reported to the WHO increasing 8-fold over the last 20 years [1]. The most significantly affected world region is The Americas reporting 3,167,542 dengue cases, 181,477 chikungunya cases and 35,914 Zika cases in 2019 alone [2–5].

In the absence of effective vaccines for dengue, Zika and chikungunya, disease control still relies on controlling mosquito vectors. Currently, this involves the use of insecticides including DDT, pyrethroids and organophosphates such as temephos, an approach that has not changed in over five decades of vector control programmes [6]. Temephos is one of the most used larvicides worldwide due to its ease of use, cost efficacy and specificity towards the larval stages of mosquitoes [7, 8]. Its pharmacological activities are related to the irreversible inhibition by phosphorylation of acetylcholinesterase (EC 3.1.1.7) [7], a ubiquitous enzyme in Metazoan primarily expressed in the nerve endings and essential for termination of acetylcholine-mediated neurotransmission [9].

Temephos was first used as a method of controlling *Ae. aegypti* larvae in the early 1970s [6] with its continued use since leading to the development of resistance in *Ae. aegypti* in multiple regions of the world [10–20]. The mechanisms conferring resistance to organophosphates have been well studied in other important vector mosquitoes but are less well understood in *Aedes* species despite their public health relevance. Mutations in the acetylcholinesterase (AChE) gene (*ace-1*) have been reported in temephos resistant insects [21] including the malaria vector *Anopheles gambiae* and the West Nile Virus vector *Culex pipiens* [22–24]. However, mutational alleles of the AChE gene are a rare finding in *Ae. aegypti* [25, 26] due to genetic constraints [27].

A commonly reported insecticide resistance mechanism in mosquitoes is the increased expression of genes encoding for enzymes capable of metabolic detoxification of insecticides [28]. Three main enzyme families have been associated with insecticide detoxification in mosquitoes: cytochrome P450 monooxygenases (P450s), glutathione S-transferases (GST) and carboxylesterases (CE). A total of 235 detoxification genes have previously been identified in *Ae. aegypti* (26 GSTs, 160 cytochrome P450s and 49 CEs) [29]. Overexpression of enzymes in all three of these groups has been associated with temephos resistance in *Ae. aegypti* [13,17,30–32]. However, the genetic and phenotypic landscapes of insecticide resistance is wider and more complex. Insecticide resistance has also been associated with cuticular modification (through alteration of cuticular thickness or composition) [33–36] and behavioural avoidance [37]. In those reports comprehensive gene expression profiling has shown great discriminatory and quantitation power for identifying a wider range of potential genes involved in insecticide resistance responses [34–36,38].

Next generation sequencing techniques, including RNA-Seq, provide a whole transcriptome approach to the identification of resistance genes with high sensitivity and specificity. RNA-Seq is now commonly used to investigate insecticide resistance in mosquitoes of medical relevance (e.g. *An. gambiae* [39], *Ae. albopictus* [40]). In *Ae. aegypti* RNA-Seq has been utilised to characterise the gene expression changes associated with insecticide resistance developed through lab selection [41–43], however this approach has sparsely been used for *Ae. aegypti* with field derived insecticide resistance [38].

This study aimed to profile mechanisms of resistance to the larvicide temephos in natural populations of *Ae. aegypti.* The field samples originated from areas of differential arbovirus burden and incidence in Colombia. In a previous study we stratified three regions in this country with distinctive arboviral disease incidence, climatic variables and socio-economic profiles through a recent time window of 11 years [44]. In the present work, *Ae. aegypti* mosquito samples from two of those regions, Bello and Cúcuta, which had the lowest and highest strata of disease burden, respectively were analysed. The differential gene expression associated with the resistance to temephos was profiled in these two field populations of *Ae. aegypti* (field-to-field comparison) with a data triangulation against the gene expression of the lab-adapted *Ae. aegypti* reference strain New Orleans (NO).

The comparison of a field resistant population against a susceptible field population and a susceptible lab population allowed us to select for gene expression specifically related to the trait under study (resistance to temephos in circulating natural populations of *Ae. aegypti*) while minimising the background noise from genotypic distance and phenotypic drifting of field mosquito populations. The present work illustrates two angles of observations in mosquito biology under selective pressure to the larvicide temephos: firstly, the potential mechanisms of insecticide resistance itself and secondly, the further adjustments in phenotype (extrapolated from gene expression profiles) that field insecticide resistant larvae undergo when in transient exposure to the insecticide. The data presented significantly expands the hitherto known composition of the gene expression responses of *Ae aegypti* mosquitoes resistant to the larvicide temephos, with a granularity at the transcriptomic level that goes beyond detoxification genes.

## Materials and methods

### *Ae. aegypti* field collection and colonisation

*Ae. aegypti* were collected from the Colombian municipalities of Bello and Cúcuta (Fig 1). These municipalities were previously shown to be distinct in burden of *Ae. aegypti* borne disease, socioeconomic status and climate [44]. Mosquito collections took place in 2016 (Bello) and 2017 (Cúcuta) with the assistance of personnel from biology and control of infectious diseases research groups (University of Antioquia) and vector-borne disease program staff within each municipality. Immature *Ae. aegypti* were collected from deployed oviposition traps (ovitraps) and reared to adults under standard conditions at Universidad de Antioquia, Colombia. Standard rearing conditions were 28 ± 1 °C, 80 ± 5% relative humidity and a 12 h light: 12 h dark photoperiod. Reared adults were offered a blood meal and the eggs collected for establishment of colonies. Upon establishment of colonies eggs were collected and sent to Edge Hill University, UK for insecticide resistance profiling.

**Fig 1.**
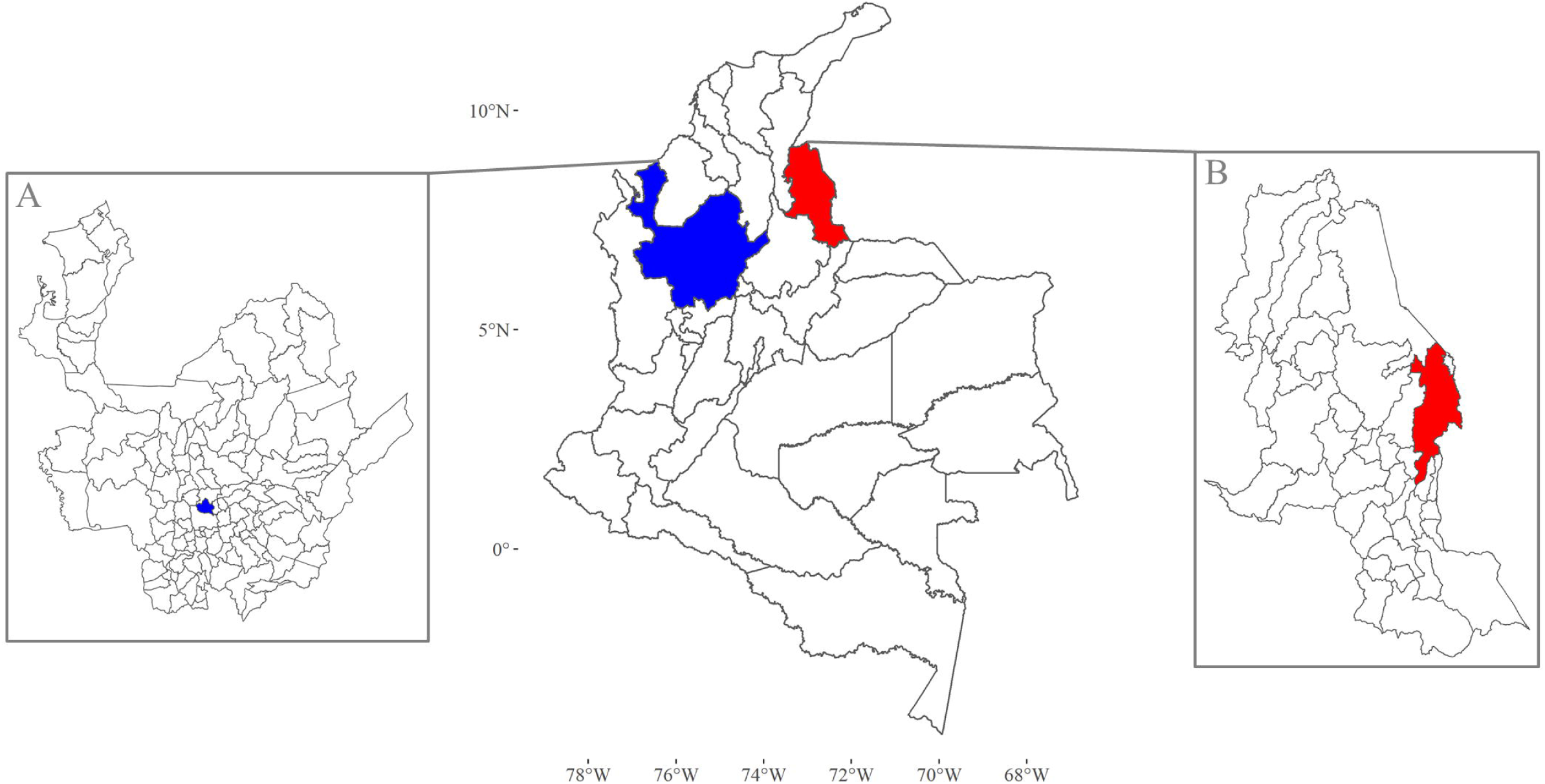
The location of the two study sites. Mosquito collections took place in these two locations, Bello and Cúcuta, within Colombia. Departments are the largest units of local government. (A) Department of Antioquia governs Bello which is denoted as a small blue area. (B) Department of Norte de Santander has as its capital Cúcuta, a city (red) to the East of this department on the border with Venezuela. Map base layers were obtained from https://data.humdata.org/dataset/colombia-administrative-boundaries-levels-0-3 covered by a Creative Commons Attribution 4.0 International (CC BY) License (https://creativecommons.org/licenses/by/4.0/legalcode). Map base layers were modified by the addition of colours.

### Larval bioassays

Larvae for use in insecticide bioassays were reared under standard conditions within Edge Hill University Vector Research Group insectaries. Standard conditions were 27°C and 70% RH with an 11-hour day/night cycle with 60-minute dawn/dusk simulation periods, using a lighting system of 4× Osram Dulux 26W 840 lights. Eggs were submerged in a hatching broth of 350 ml distilled H_2_O (dH_2_O), 0.125 g nutrient broth (Sigma-Aldrich, Darmstadt, Germany) and 0.025 g brewer’s yeast (Holland & Barrett, Ormskirk, UK) for 48 hours [45]. Larvae were fed ground fish food (AQUARIAN® advanced nutrition) and raised until third to fourth instar. Larval bioassays were conducted following WHO standard test procedures [46]. Preliminary testing was conducted to identify the activity range of temephos to larvae from each of the study municipalities and a susceptible laboratory strain (New Orleans). The activity ranges, yielding mortality of 10 -95%, for each *Ae. aegypti* population as identified by preliminary testing are displayed in Table 1. At least four replicates, each consisting of 20 third to fourth instar larvae, were conducted for each temephos concentration. Fresh insecticide solutions were made for each replicate using temephos (93.7%; Pestanal®, Sigma-Aldrich Darmstadt, Germany) and acetone (≥ 99.9%; Sigma-Aldrich, Darmstadt, Germany). Bioassays were conducted inside the insectaries under standard conditions and mortality recorded after a 24-hour exposure period. Following WHO guidelines, moribund larvae were counted as dead. Controls were exposed to the acetone solvent only.

**Table 1.**
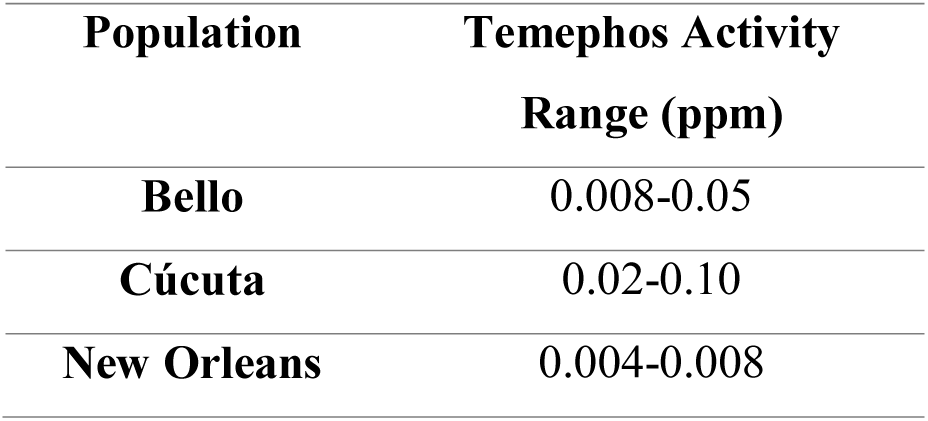
Temephos activity range yielding between 10-95% mortality to each mosquito population. Results of preliminary testing to identify the activity range of temephos to each mosquito population. Activity ranges displayed in parts per million (ppm).

### RNA extraction and cDNA synthesis

#### Aedes aegypti sample groups

Following the larval bioassays, field samples were categorised as resistant (Cúcuta) or susceptible (Bello). In the latter category were also samples from the lab adapted reference strain New Orleans. The Cúcuta temephos resistant samples were further divided into two groups: one exposed to temephos (for 24 hours) immediately before sampling for RNA extraction, and a control group of no temephos exposure (unexposed) (Fig 2). For each group (field susceptible (FS), lab susceptible (LS), field resistant exposed (FRE) and field resistant unexposed (FRU)) RNA extractions were carried out from four different larvae batches considered here as biological replicates.

**Fig 2.**
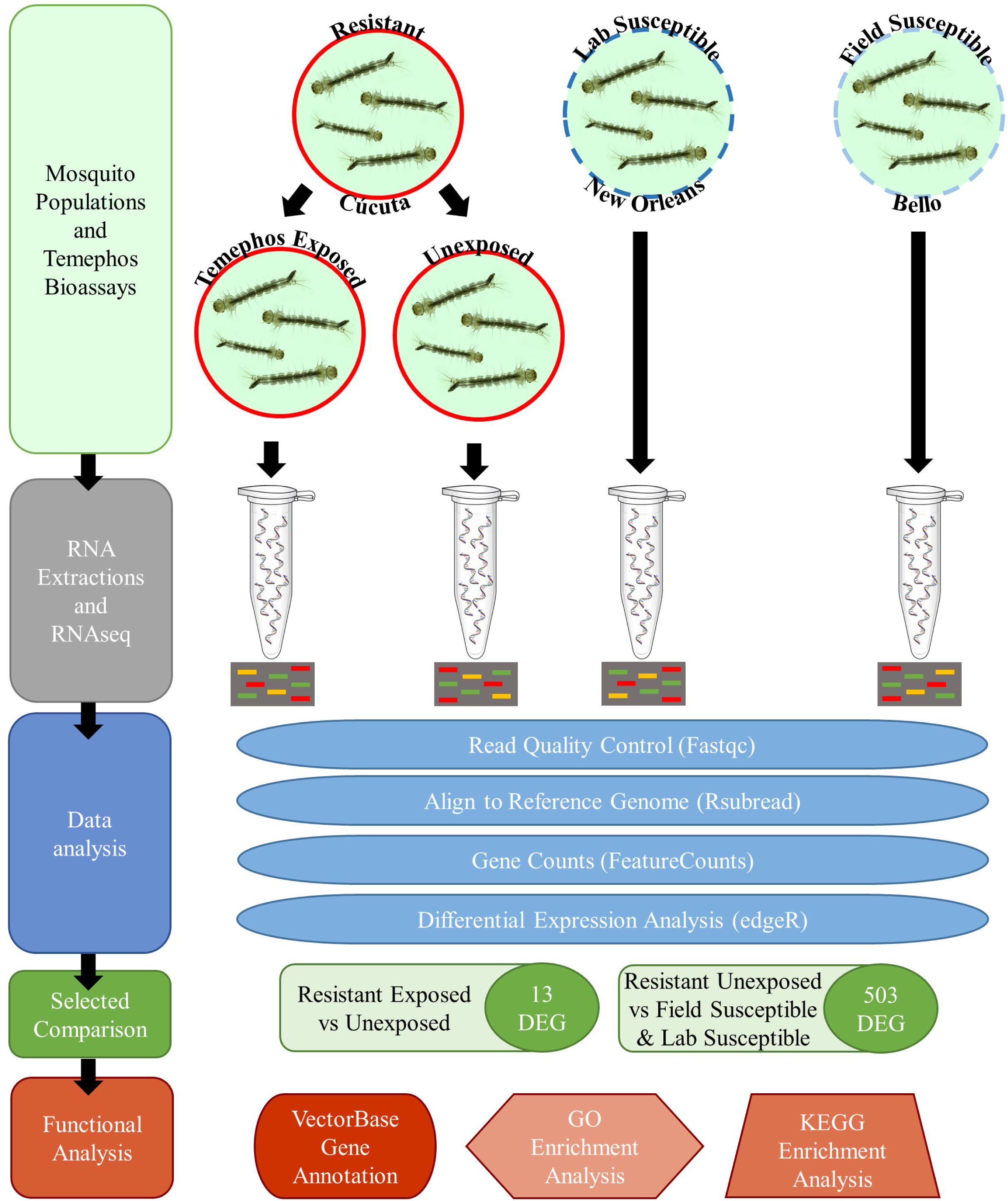
Block diagram of our experimental approach. This study concerned the larval stages of the mosquito *Ae. aegypti*. Field samples denoted as Resistant originated from the Cúcuta, Colombia population. The Susceptible samples had dual origin: Field samples from Bello, Colombia (Field Susceptible) and the New Orleans reference lab strain (Lab Susceptible). The total RNA sequenced and mapped (Data analysis) originated from four different experiments (biological replicates) from each population. The gene expression levels of the Resistant samples compared against the Lab Susceptible and Field susceptible had at least 503 differentially expressed genes (DEG). The Resistant samples of larvae transiently exposed to temephos had 13 DEGs in comparison to the unexposed larvae. The functional annotation for the DEG sets was carried with several different repositories: VectorBase, Gene Ontology (GO) and KEGG Enrichment Analysis.

#### Standard rearing of *Aedes aegypti* for RNA extraction

*Ae. aegypti* were reared to fourth instar larvae following a standard rearing protocol and under standard conditions within Edge Hill University Vector Research Group insectaries. Standard conditions were 27°C and 70% RH with an 11-hour day/night cycle with 60-minute dawn/dusk simulation periods, using a lighting system of 4× Osram Dulux 26W 840 lights. Eggs were submerged in a hatching broth of 350 ml dH_2_O, 0.125 g nutrient broth (Sigma-Aldrich, Dorset, UK) and 0.025 g brewer’s yeast (Holland & Barrett, Ormskirk, UK) for 48 hours [45]. Once hatched, larvae were reared at a density of one larva/ml in dH_2_O and fed ground fish food (AQUARIAN® advanced nutrition) at increasing quantities per day (day 3 = 0.08 mg/larva, day 4 = 0.16 mg/larva, day 5 = 0.31 mg/larva) [47]. Six days after egg submersion larvae were subjected to an insecticide bioassay in batches of 25 larvae in 100 ml dH_2_O, for 24 hours [46].

#### Temephos exposure assays for resistant larvae

Larvae in the resistant exposed group were exposed to temephos at the LC_50_ of 0.06 ppm (Fig 2). Larvae in all unexposed groups were exposed to the equivalent volume of acetone (the solvent used in temephos solutions). After the 24 h bioassay larvae were taken for RNA extraction. For each experimental group there were four independent replicates, conducted using eggs from different batches and rearing and extraction conducted on different days. Egg submersion, feeding, bioassays and RNA extraction on all replicates were all conducted at the same times of day.

#### RNA extraction

Larvae were homogenized using QIAshredders (Qiagen, Manchester, UK) then RNA extracted using PicoPure® RNA Isolation Kit (Arcturus Bioscience, Mountain View, USA) following the manufacturers’ protocols. RNA was extracted from a total of 20 individual larvae per biological replicate, with four larvae per column and the total RNA then pooled. RNA quality and quantity were assessed using an Agilent 2100 bioanalyzer. The temephos exposed population had 12-16 larvae per replicate due to mortality during bioassays.

#### Library preparation and sequencing

Library preparation and sequencing was conducted at Polo d’Innovazione di Genomica, Genetica e Biologia, Italy. Libraries were prepared following the QIAseq^TM^ Stranded mRNA Select Kit Handbook (June 2019) for Illumina Paired-End Indexed Sequencing [48]. Libraries were validated using the Fragment Analyzer High Sensitivity Small Fragment method to assess size distribution and quantified using a Qubit® 3.0 Fluorometer. Indexed DNA libraries were all normalized to 4 nM, pooled in equal volumes, and then loaded at a concentration of 360 pM onto an Illumina Novaseq 6000 S1 flowcell, with 1% of Phix control. The samples were sequenced using the Novaseq 6000 standard workflow with 2 x 150 bp pair end run. The experimental design for this study is outlined in Fig 2. The raw reads obtained through RNA-Seq are deposited in NCBI’s sequenced read archive (Accession PRJNA730411).

### Data analysis

#### Bioassay data analysis

Larval 50% and 95% lethal concentrations (LC_50_ and LC_95_) and their 95% confidence limits (p < 0.05) were calculated using probit analysis according to Finney (1947) [49] using the LC_probit function in the R ecotox package (version 1.4.0) [50]. Abbot’s correction [51] was not applied due to the control mortality never exceeding 10%. Resistance ratios (RR_50_ and RR_90_) were calculated to assess temephos resistance by comparison of LC_50_ and LC_90_ for *Ae. aegypti* from each field location to those of the susceptible laboratory strain (New Orleans). Resistance ratios were defined as susceptible (<5-fold), moderate resistance (5 - 10-fold) and high resistance (>10-fold) following WHO guidelines [46]. Statistical analyses of bioassay data were conducted using R statistical software (version 3.6.1) [52].

#### RNA-Seq data quality control, mapping and differential gene expression

Analyses of RNA-Seq data were conducted using the Linux command line (Ubuntu 18.04) and R statistical software (version 4.0.3) [52]. Sequence read quality was assessed using FastQC (version 0.11.3). Reads with quality scores less than 30 and lengths less than 50 bp were trimmed using cutadapt (version 2.10) with Python (version 3.8.3). Read quality was then reassessed using FastQC to ensure only high-quality reads remained. Cleaned reads were mapped to the *Ae. aegypti* LVP_AGWG reference genome (version AaegL5, GenBank: NIGP00000000.1) using Rsubread (version 2.2.6) [53]. The resultant BAM files were sorted and indexed using samtools (version 1.11). Alignment quality metrics from Rsubread were visualized using R’s plotting function. Gene count tables were generated using Rsubreads. Read counts were normalized using the trimmed mean of M values method [54] in edgeR (Version 3.30.3) which accounts for library size and expression bias in RNA-Seq datasets [55]. Differential gene expression was then calculated using a Quasi-likelihood negative binomial generalized log linear model (edgeR). Quasi-likelihood error family was selected due to its ability to account for uncertainty in dispersion. Counts per million (CPM) were calculated in edgeR and reads per kilobase million (RPKM) were calculated in edgeR using transcript lengths obtained from Enseml Metazoa (LVP AGWG (aalvpagwg_eg_gene) dataset) using biomaRt (version 2.44.4). Transcripts with fold-change >2 and FDR < 0.05 were selected for gene ontology and KEGG pathway enrichment analyses.

#### Gene ontology and KEGG pathway enrichment analyses

Gene ontology (GO) category assignments were obtained from Ensembl Metazoa using biomaRt (version 2.44.4) [56] and KEGG pathway assignments from Kyoto Encyclopedia of Genes and Genomes using KEGGREST (1.28.0) [57]. GO and KEGG enrichment analyses were conducted using GOseq (version 1.40.0), which allows for correction of biases arising from the variable transcript lengths in RNA-Seq data [58]. Enrichment scores were calculated using the Wallenius method within GOseq. P-values were then corrected for multiple testing using the Benjamini–Hochberg method in the p.adjust function [52]. GO categories and KEGG pathways with corrected p values <0.05 were considered significantly enriched. Enrichment percentage was calculated as a ratio of the number of differentially expressed genes within each category to the total number of genes within that category.

## Results

### Temephos susceptibility of *Ae. aegypti* field isolates and the reference strain

The lethal concentrations (LC_50_) of temephos were 0.019 *ppm* (95% Cl 0.016-0.029) and 0.060 *ppm* (Cl 95% 0.052-0.070) for Bello and Cúcuta, respectively. This corresponded to resistance ratios of 2.6 and 8.0 when compared to the New Orleans susceptible laboratory strain (LC_50_ = 0.008; Cl 95% 0.006-0.012). The LC_95_ for Bello was 0.055 (Cl 95% 0.034-0.292), 2.8-fold higher than New Orleans (LC_95_ = 0.020; Cl 95% 0.012-0.21). Cúcuta had a LC_95_ of 0.182, 9-fold higher than New Orleans (Table 2). Following the WHO guidelines [46] larvae from Bello were considered susceptible to temephos and denoted as the field susceptible (FS) population whist the larvae from Cúcuta were resistant to this larvicide and denoted as the field resistant (FR) population (Table 2). New Orleans is referred to as the lab susceptible (LS) population.

**Table 2:**
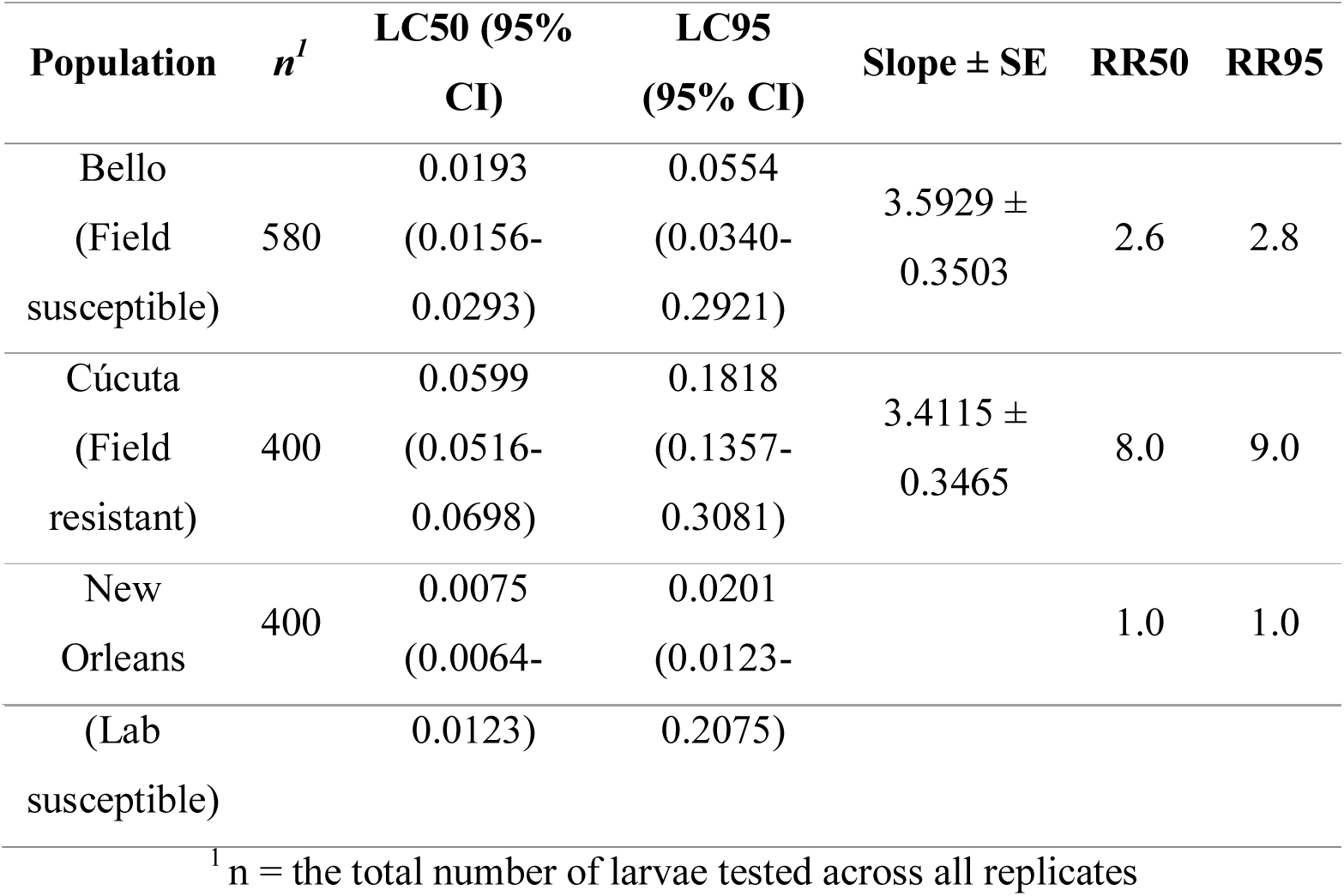
LC_50_ and LC_95_ of Bello, Cúcuta and New Orleans Ae. aegypti larvae to temephos. Temephos bioassays showing LC50, LC95 and their 95% confidence limits, calculated using probit analysis. Resistance ratios (RR) calculated as a ratio of the lethal concentration (LC50 and LC95) of each population compared to the lab susceptible (New Orleans) strain. Bello and New Orleans are both susceptible to temephos whilst larvae from Cúcuta are resistant. SE = standard error.

### RNA-Seq mapping summary

Three different populations of *Ae. aegypti* larvae were profiled using RNA-Seq: temephos field resistant (FR), field susceptible (FS) and the reference lab susceptible strain New Orleans (LS). The FR population was also split in two further groups as either exposed or not (unexposed) to temephos to determine further expression changes associated with insecticide exposure in already resistant populations (Fig 2). Mosquito larvae from each of these four populations were grown in four different batches, one to four weeks apart and were considered here four biological replicas. This generated a total of 16 sequenced samples.

Extracted total RNA was used to generate the Illumina RNA-Seq libraries that produced 65.7 – 120 million reads per sample with quality scores >30 and lengths >50 bp and 84.4% and 85.6% of those reads in each sample were successfully aligned to the reference genome (Table 3 and Materials and Methods). Using the most current gene model available for *Ae. aegypti,* LVP_AGWG reference genome which contains 19,381 open reading frames (version AaegL5, GenBank: NIGP00000000.1) the number of genes with successfully aligned reads ranged from 15,704 and 16,607 genes across all samples, corresponding to 81 – 86% of the total genes in the reference genome.

**Table 3:**
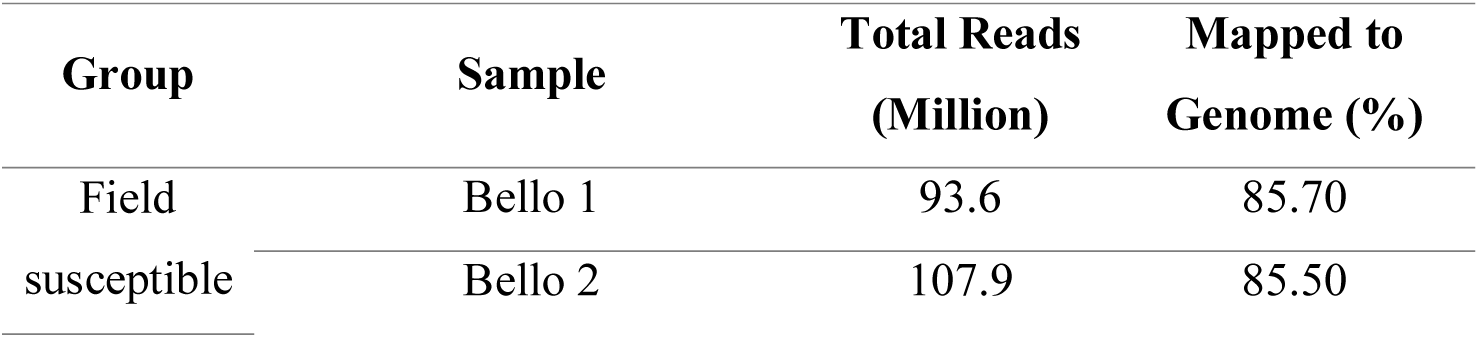
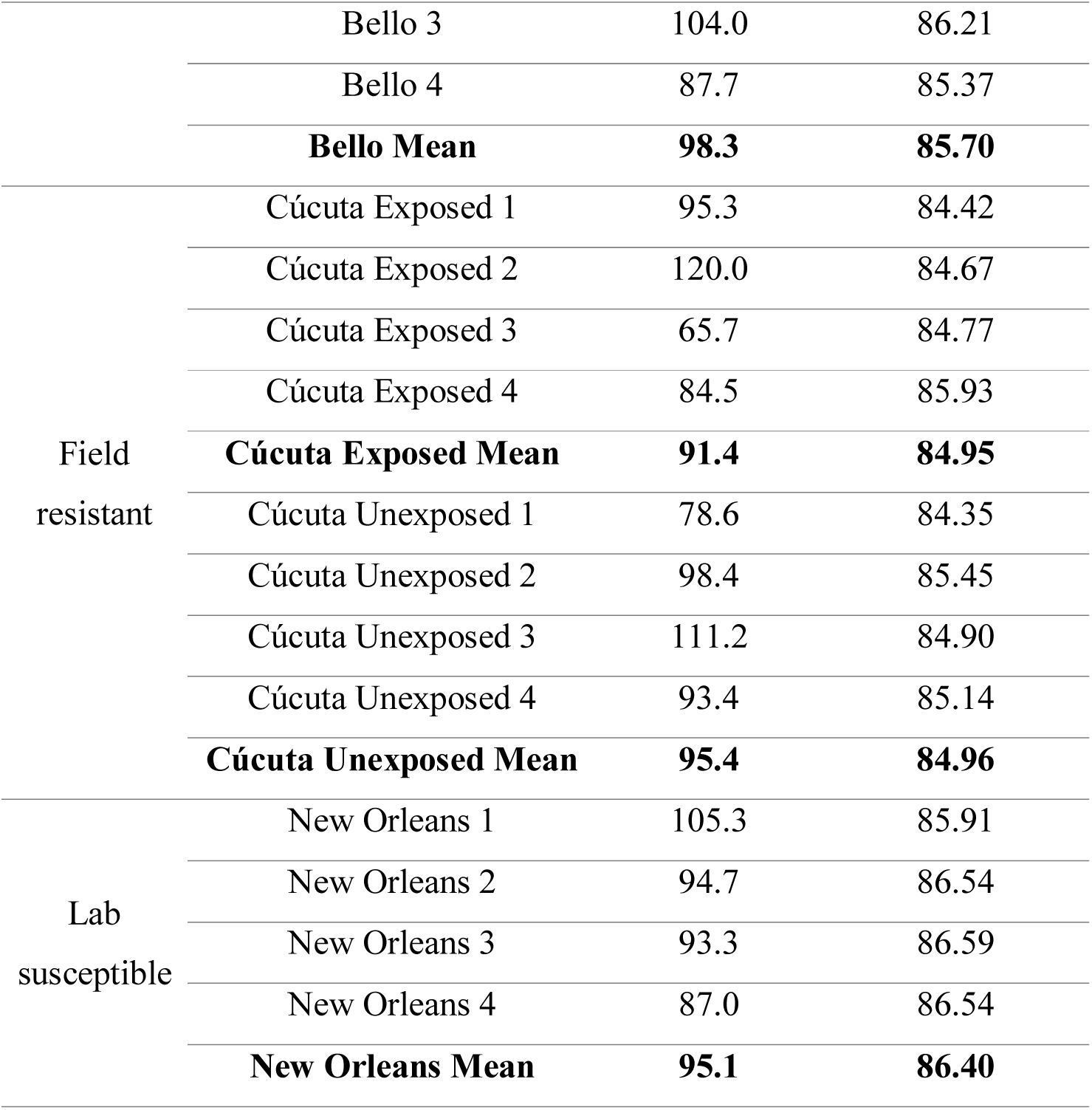
RNA-Seq sequencing data summary. The total number of obtained reads after quality control and the percentage of reads that mapped to the *Aedes aegypti* reference genome. Quality control removed reads with quality <30 and lengths <50bp.

### Overview of differential gene expression

Both field samples, susceptible and resistant (FS, FR) were equally and evidently distant, by RPKM number, to the samples of the lab strain New Orleans (LS) (Fig 3). Crucially the temephos susceptible samples did not cluster together nor did the two field samples (Fig 3). Field and lab strain samples were similarly distant in terms of differentially expressed (DE) transcripts: FS vs LS = 5324 DE transcripts (Fig 4D and 4E), FR vs LS = 5579 (Fig 4B and 4E). However, when comparing field samples (resistant and susceptible) the number of DE transcripts visibly lowered by four-fold to 1,454 (Fig 4A and 4E). Therefore, the common practice in gene expression studies of comparing mosquito field samples to lab strains (e.g. [17,30,59,60]), would have generated an approximately 4-fold overestimation in the number of DE transcripts detected in the field resistant samples.

**Fig 3.**
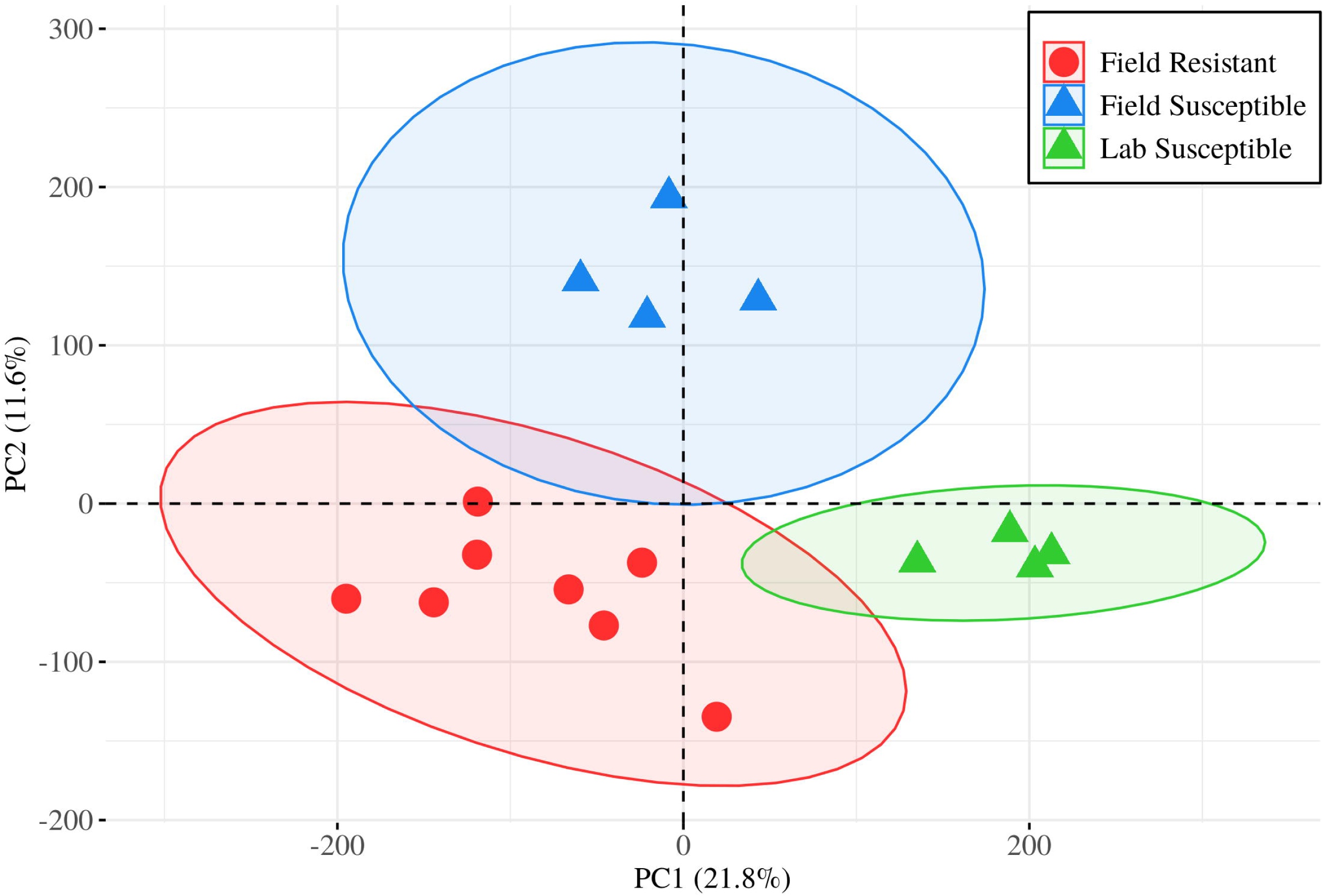
Distribution of the transcriptomic profiles by sample groups. The multidimensionality of the RPKM values calculated for each mapped transcript per sample was reduced by principal components analysis (PCA). The field samples were seen linearly distant from each other across one component whist the reference lab strain NO cluster (Lab Susceptible) separated from both field samples. The orthogonal dispersion of these samples allowed for the triangulation of the data as described in the main text.

**Fig 4.**
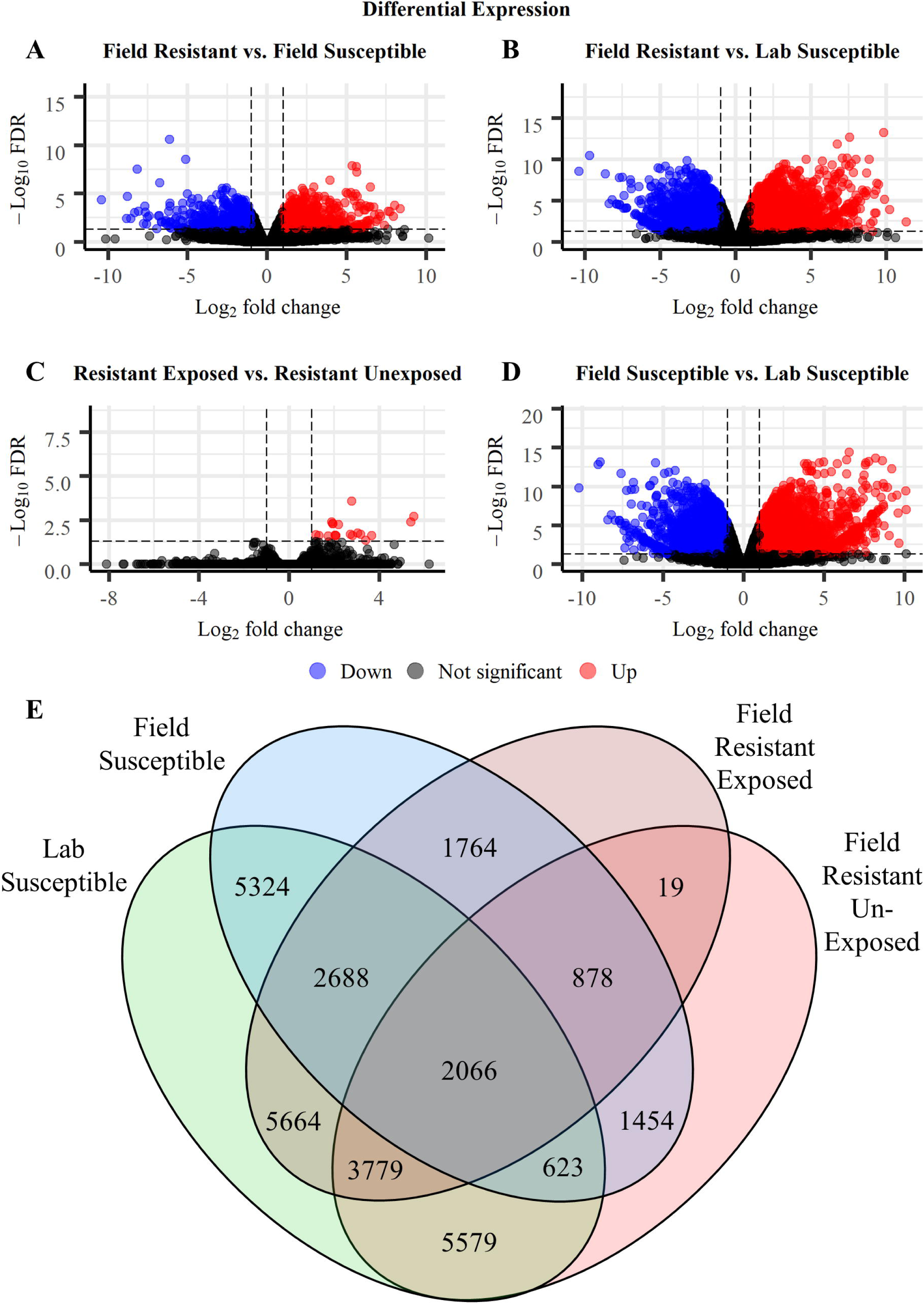
Differential gene expression in the samples from field susceptible, lab susceptible, and field resistant populations. (A-D) Differential expression of all transcripts including those over expressed (red), under expressed (blue) and with no significant differential expression (black) in the field resistant unexposed population compared to the field susceptible population (A) and susceptible lab population (B), the resistant temephos exposed population when compared to the resistant unexposed population (C) and in the field susceptible population when compared to the lab susceptible population (D). (E) The number of transcripts significantly differentially expressed (FC >2, FDR <0.05) between each of the experimental groups. The number of differentially expressed transcripts shown here (E) include both significantly over and under expressed transcripts. The comparison groups and sample notation as detailed in Fig 2.

We also sought to investigate the gene expression changes in insecticide resistant larvae under transient exposure to insecticide. There were only 19 transcripts significantly differentially expressed in larvae within the resistant population which were exposed to temephos when compared to unexposed larvae from the same population (Fig 4E). All 19 of those transcripts were overexpressed in the exposed group with no significant down regulated gene expression detected (Fig 4C).

We addressed the issue of potential misrepresentation of gene expression metrics by triangulating both, the differentially expressed gene (DEG) sets and the RPKM counts between the field resistant (FR) samples against the field susceptible (FS) as well as the lab susceptible (LS) samples. The DEG set obtained contained transcripts which were found to be significantly differentially expressed with a fold change of >2 and a false-discovery rate (adjusted P value) of <0.05 in both comparisons. Under this threshold, a total of 623 (down from 1,454 transcripts in only the field-to-field comparison) transcripts covering 503 genes, were found differentially expressed in the field resistant population when compared to both the field and lab susceptible populations (Fig 4E). This set of 503 genes comprised 301 overexpressed genes and 202 under expressed genes (S1 Table). Of the 301 significantly overexpressed genes 239 were found in the category of protein coding genes: 88 annotated and 151 hypothetical genes. In the significantly under expressed gene set 166 were protein coding with 75 annotated and 91 hypothetical genes (S1 Table). The significant differentially expressed genes also included 55 overexpressed genes encoding for long non-coding RNA (lncRNA) and 30 under expressed lncRNA genes in the temephos resistant larvae (S1 Table).

### Gene ontology and KEGG pathway enrichment

All 623 transcripts with fold-change >2 and FDR < 0.05 were selected for gene ontology (GO) and KEGG pathway enrichment analyses. GO categories and KEGG pathways with corrected p values (FDR) < 0.05 were considered significantly enriched. Gene ontology and KEGG enrichment analyses conducted on the 301 significantly over expressed genes identified eight significantly enriched GO categories; one involved with biological processes, oxidation-reduction processes (GO:0055114), and seven associated with molecular functions (GO:0045735, GO:0016705, GO:0004100, GO:0004022, GO:0005506, GO:0016491, GO:0047938) (Fig 5). KEGG enrichment analysis identified two significantly enriched KEGG pathways in the over expressed genes; insect hormone biosynthesis (path:00981) and ubiquinone and other terpenoid-quinone biosynthesis (path:00130) (Fig 5).

**Fig 5:**
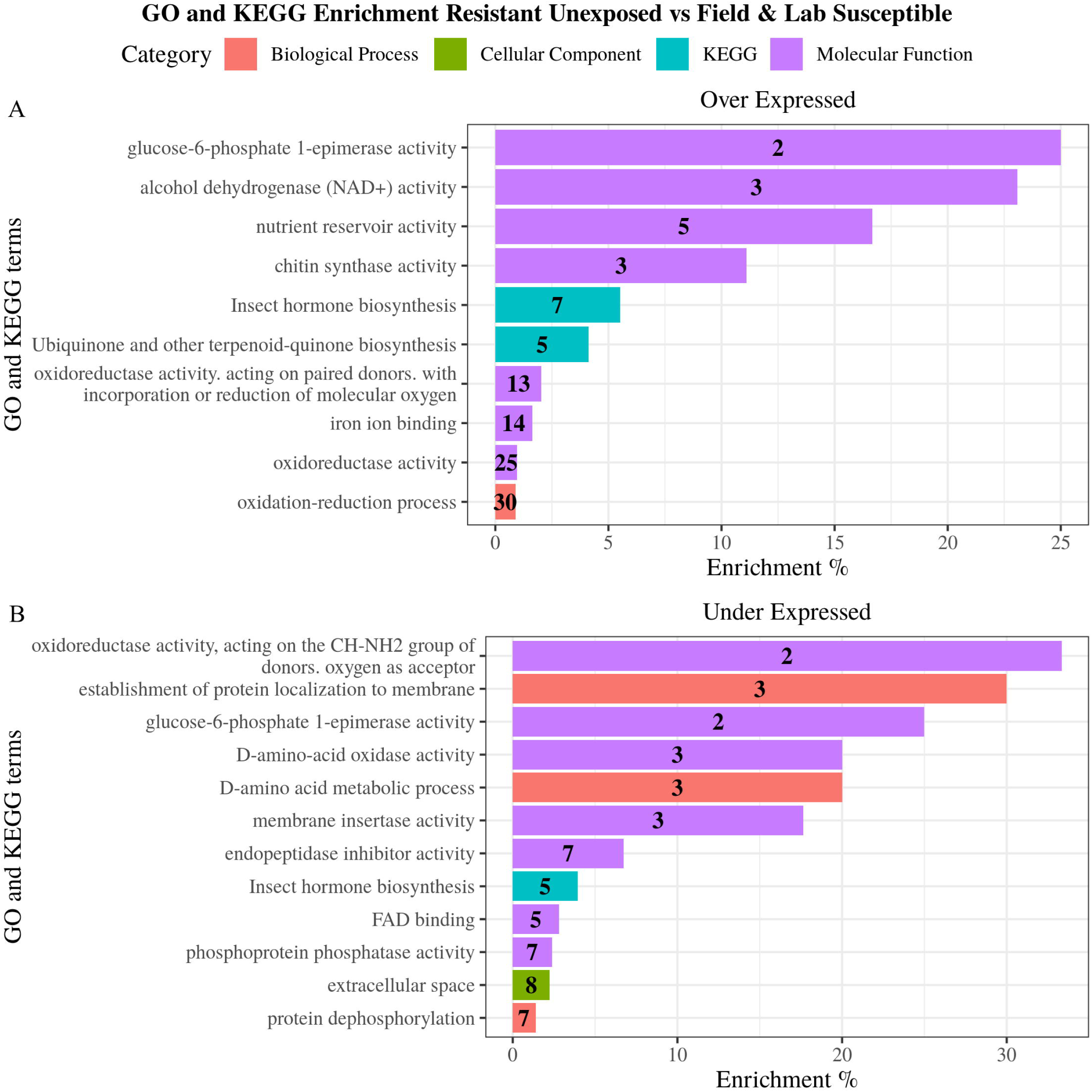
GO terms and KEGG pathways enriched in the resistant population when compared to both field and lab susceptible populations. GO terms and KEGG pathways found to be significantly enriched (p <0.05) following Benjamini Hochberg correction in the significantly over (A) and under (B) expressed transcripts. Enrichment percentage was calculated as the number of differentially expressed transcripts in each category/pathway divided by the total number of transcripts in the same category/pathway. Number in bars indicate the number of differentially expressed transcripts in each category.

GO and KEGG enrichment analysis were also conducted on the 202 significantly under expressed genes identifying 12 significantly enriched GO terms and one KEGG pathway (path:00981) (Fig 5). The enriched GO terms include three terms involved in biological processes (GO:0090150, GO:0046416, GO:0006470), one involved with cellular components (GO:0005615) and seven associated with molecular functions (GO:0004866, GO:0004721, GO:0003884, GO:0032977, GO:0047938, GO:0016641, GO:0071949) (Fig 5).

### The overexpressed transcriptome of field temephos resistant *Aedes aegypti* larvae

The transcriptomic overview provided by the GO and KEGG enrichment models was interrogated by quantifying the represented genes with CPM and RPKM metrics. The expression profiles of the 88 annotated overexpressed protein coding genes in the resistant population were visualised using heatmaps of gene expression as log_2_ values of counts per million (CPM) (Fig 6) as well as bar plots of reads per kilobase million (RPKM) values of the represented genes (Fig 7). The former allowed the visualisation of the data’s granularity by comparing gene expression across all 16 samples individually rather than just across groups (Fig 6). Variation in expression between samples from the same experimental group can be seen across all genes (Fig 6) highlighting the importance of biological replication in gene expression experimentation. The heatmap also showed the importance of using field susceptible populations in addition to lab susceptible comparator populations. Differences in gene expression (e.g., CYP6B1 and PGRPLA) can be overrepresented when comparing expression profiles between field and lab populations (here FR and LS) rather than between field to field (e.g., FR and FS). Differences in gene expression were also visualised using RPKM bar plots which enable ranking of genes specifically overexpressed in resistant samples, the majority (151 of the 239 protein coding genes) of those did not have a functional annotation in the current repository for VectorBase (Fig 7, S1 Table).

**Fig 6:**
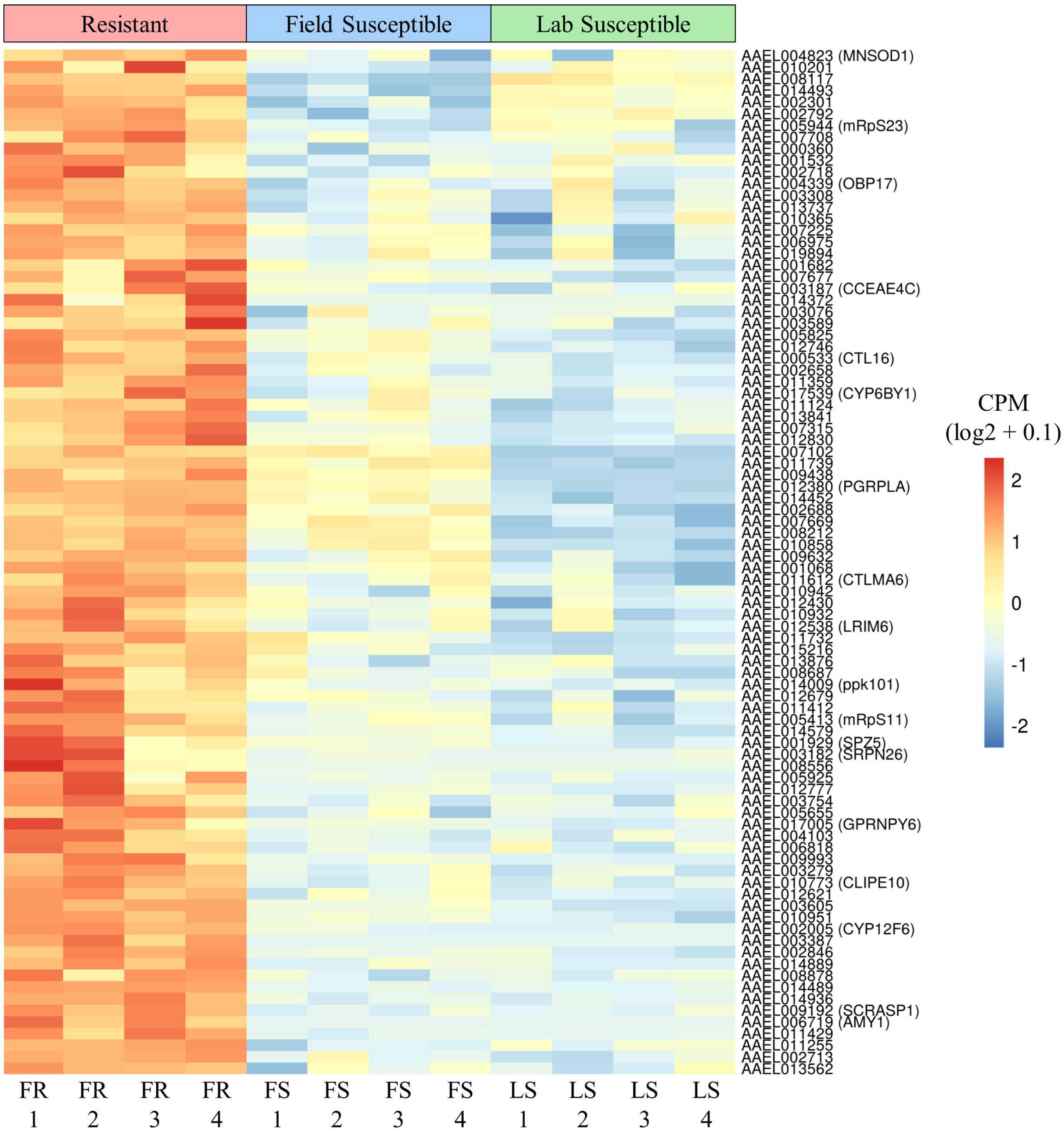
Comparison of gene expression of significantly over expressed protein coding annotated genes in the field resistant, field susceptible and laboratory susceptible populations. Comparison of the FR population to both FS and LS populations identified a total of 88 protein coding VectorBase annotated genes with significant over expression (FC >2, FDR <0.05). The expression levels were displayed as counts per million (CPM) which is the number of counts per gene following normalisation. CPM values were calculated for each gene by taking the mean CPM of each transcript within that gene. Gene expression was displayed per sample, rather than per experimental group, allowing for visualisation of granularity between samples. The gene expression was scaled by row to allow comparison between samples rather than between genes. Variability in expression between samples from the same experimental groups can be seen across all genes, highlighting the importance of biological replication. There was a larger difference in expression between the FR and LS than between the FR and FS for many genes including CYP6B1 and AAEL007102, demonstrating the importance of using multiple susceptible comparator strains to reduce over estimation of DGE.

**Fig 7:**
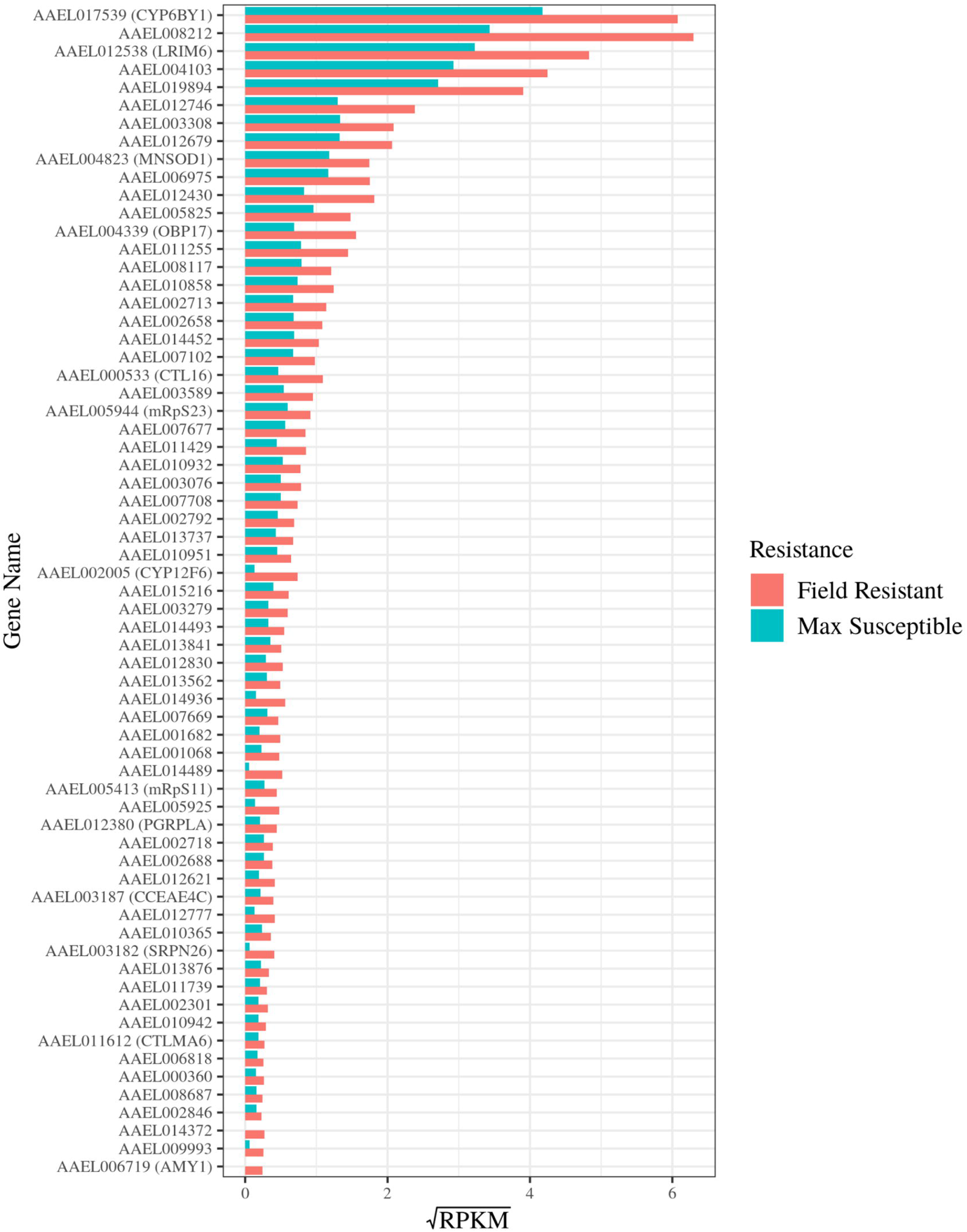
Differences in gene expression between resistant and susceptible populations of a set of significantly over expressed, protein coding genes with VectorBase annotation. Comparison of the FR population to both FS and LS populations identified a total of 88 protein coding VectorBase annotated genes with significant over expression (FC >2, FDR <0.05). Gene expression was displayed here as reads per kilobase million (RPKM) in the FR population (red) and both susceptible populations (turquoise). RPKM values were calculated for each gene by taking the mean RPKM of each transcript within that gene. The susceptible RPKMs (max susceptible) represent the maximum RPKM for each gene in both FS and LS populations. The RPKM values were square root transformed here to optimise the visualisation of a vast range of values (0.02 to 39.11). Genes with average RPKM across groups of below 0.02 (23 genes) were not included in the bar plot for visualisation purposes but are included in S1 Table. Mean RPKM values per resistance status allow for comparison of expression between genes as well as between groups.

The over expressed annotated protein coding genes included detoxification enzymes; two cytochrome P450s (*CYP12F6 -* AAEL002005 and *CYP6BY1 -* AAEL017539), a carboxy/cholinesterase (*CCEAE4C* - AAEL003187), a glutathione S-transferase (AAEL006818), two glucosyl/glucuronosyl transferases (AAEL002688 and AAEL003076) and an aldehyde oxidase (AAEL014493). The cuticular biosynthesis enzyme chitin synthase (AAEL002718) and the digestive enzymes, putative trypsin genes, AAEL007102, AAEL014579, and AAEL003308, were also present.

Other over expressed genes included the hydrocarbon biosynthesis pathway enzyme acetyl-CoA dehydrogenase (AAEL014452) [61], glutamate decarboxylase (AAEL010951) which catalyses biosynthesis of GABA through glutamate decarboxylation [62], sarcosine dehydrogenase (AAEL014936), a mitochondrial glycine synthesising enzyme [63], leucine aminopeptidase (AAEL006975), a proteolytic enzyme that hydrolyses amino acids with roles in toxin biosynthesis [64], a manganese-iron (Mn-Fe) superoxide dismutase (*MNSOD1 -* AAEL004823), a mitochondrial antioxidant associated with increased life span in insects [65, 66] and two mannose-binding C-Type Lectins (CTLs) AAEL011612 and AAEL000533, ubiquitous proteins in multicellular organisms that provide the pattern recognition required for the initial phase of an immune response [67, 68].

### The under expressed transcriptome of field resistant *Aedes aegypti* larvae

The transcript profiles of the 75 annotated protein coding genes significantly under expressed genes in the resistant population were also visualised in heatmaps of gene expression (log_2_ values of CPM) (Fig 8) as well as bar plots of RPKMs (square root values) of the represented genes (Fig 9). The former showing variation between the 16 samples and the latter displaying variation between genes and between resistance status.

**Fig 8:**
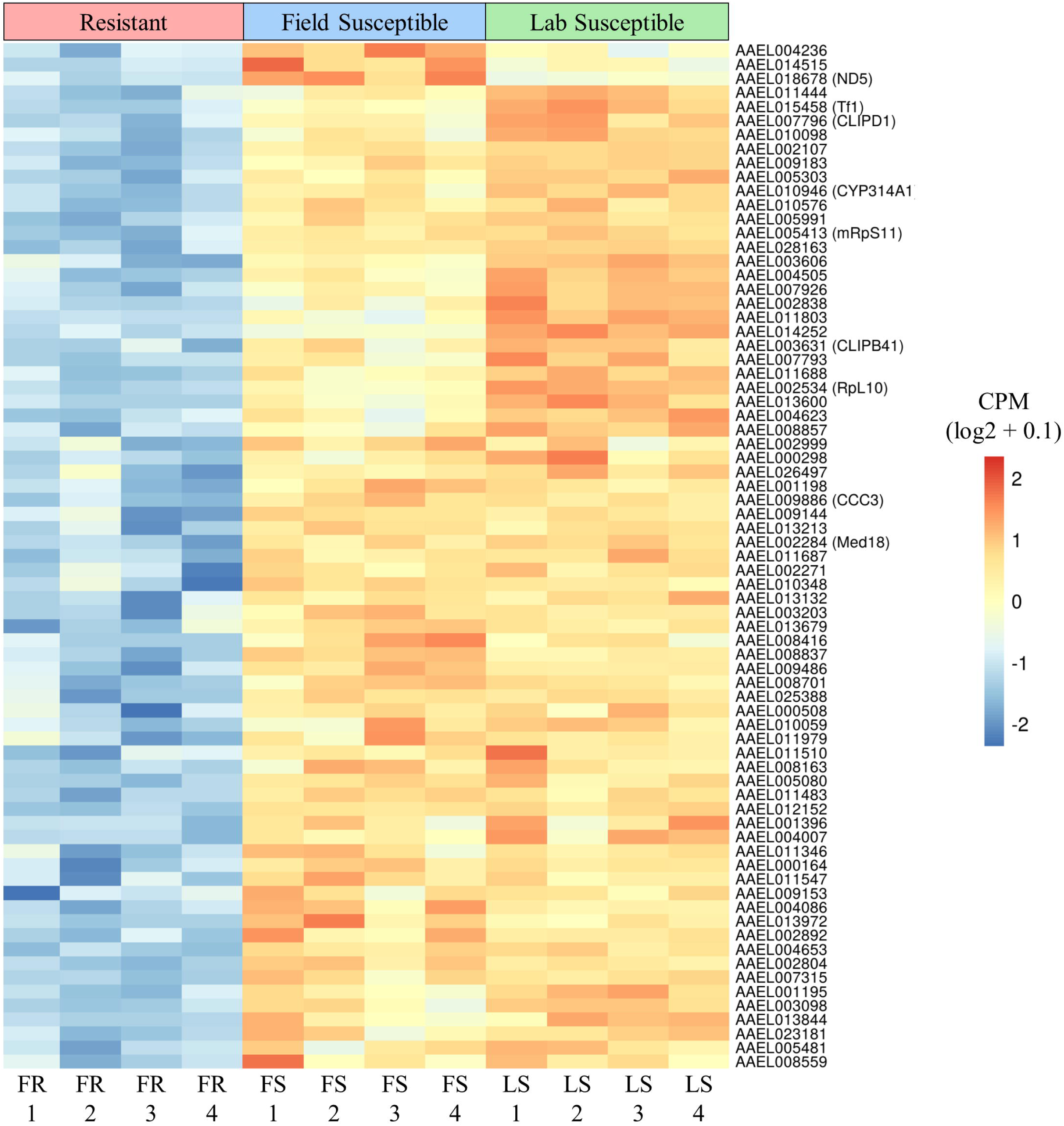
Comparison of gene expression of significantly under expressed protein coding annotated genes in the field resistant, field susceptible and laboratory susceptible populations. Comparison of the FR population to both FS and LS populations identified a total of 76 protein coding VectorBase annotated genes with significant under expression (FC >2, FDR <0.05). The expression was displayed here as counts per million (CPM) which is the number of counts per gene following normalisation. CPM values were calculated for each gene by taking the mean CPM of each transcript within that gene. Expression was displayed per sample, rather than per experimental group, allowing for visualisation of granularity between samples. The expression was scaled by row to allow comparison between samples rather than between genes. Variability in expression between samples from the same experimental groups can be seen across all genes, highlighting the importance of biological replication. There was a larger difference in expression between the FR and LS than between the FR and FS for many genes including CLIPB41 and RpL10, demonstrating the importance of using multiple susceptible comparator strains to reduce over estimation of DGE.

**Fig 9:**
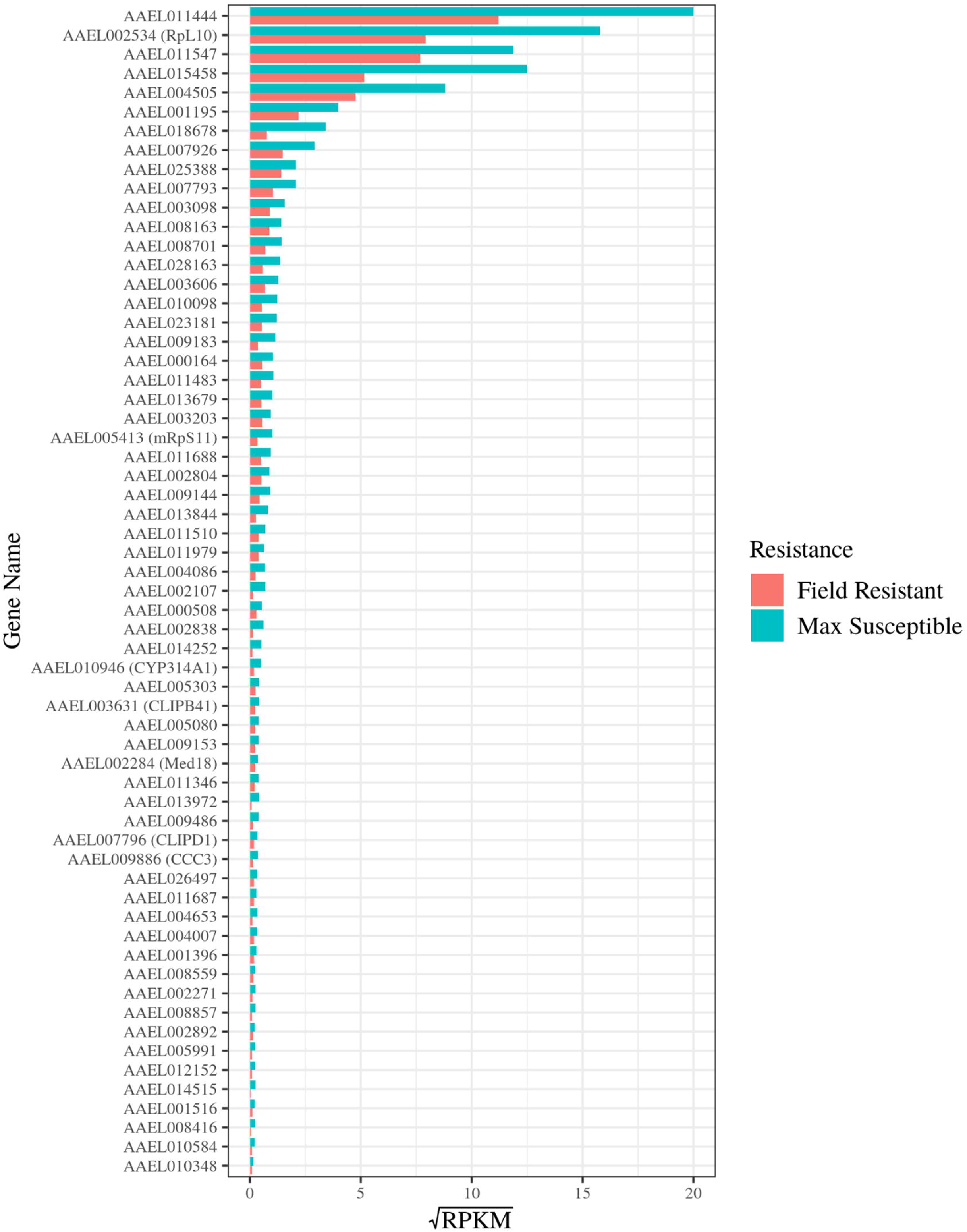
Differences in gene expression between resistant and susceptible populations of a set of significantly under expressed, protein coding genes with VectorBase annotation. Comparison of the FR population to both FS and LS populations identified a total of 76 protein coding VectorBase annotated genes with significant under expression (FC >2, FDR <0.05). Gene expression was displayed here as reads per kilobase million (RPKM) in the FR population (red) and both susceptible populations (turquoise). RPKM values were calculated for each gene by taking the mean RPKM of each transcript within that gene. The susceptible RPKMs (max susceptible) represent the maximum RPKM for each gene in both FS and LS populations. The RPKM values were square root transformed here to optimise the visualisation of a vast range of values (0.02 to 400). Genes with average RPKM across groups of below 0.02 (14 genes) were not included in the bar plot for visualisation purposes but are included in S1 Table. Mean RPKM values per resistance status allow for comparison of expression between genes as well as between groups.

Genes encoding detoxification enzymes were also presented in this set of under expressed genes. Those included the cytochrome P450 *CYP314A1* (AAEL010946) and a glucosyl/glucuronosyl transferase (AAEL003098). A cytochrome oxidase biogenesis protein (oxa1 mitochondrial - AAEL009183), essential for full expression of cytochrome c oxidase was also under expressed. Other genes significantly under expressed in the resistant population include a putative pupal cuticle protein (AAEL011444) and transferrin (*Tf1* - AAEL015458) a regulator of iron metabolism with roles in mosquito innate immunity [69]. The mdg4-binding protein ortholog gene in *Ae. aegypti* (AAEL010576: Modifier of mdg4 [Mod(mdg4)]), responsible for chromosome remodelling was also represented in this group of under expressed genes.

The expression of several ion and solute membrane transporters were also down regulated. These included the sodium-coupled cation-chloride cotransporter AAEL009886 (aeCC3), the sodium/chloride dependent amino acid transporter AAEL000298, the sodium/solute symporter AAEL001198, and the sugar transporter AAEL010348. In the group of under expressed genes were also 30 lncRNA genes in the temephos resistant larvae (S1 Table).

### Gene expression profile of temephos exposed larvae from the resistant population

Gene expression in the field resistant population following the controlled exposure to temephos was compared with gene expression of samples from the same population without insecticide exposure. The exposed samples showed 19 significantly (FC >2, p-value <0.05) overexpressed transcripts (Fig 10A & S2 Table) in comparison to the non-exposed samples. These 19 transcripts were mapped to 13 genes (Fig 10). The products of the over expressed genes include a sodium/chloride dependent amino acid transporter (AAEL003619), an alkyl dihydroxyacetone phosphate synthase (AAEL007793), cathepsin-1 (AAEL011167), trypsin - 1 (AAEL016975) and a serine protease stubble (AAEL020367). The remaining eight overexpressed genes had uncharacterised products in *Ae. aegypti* (S2 Table).

**Fig 10:**
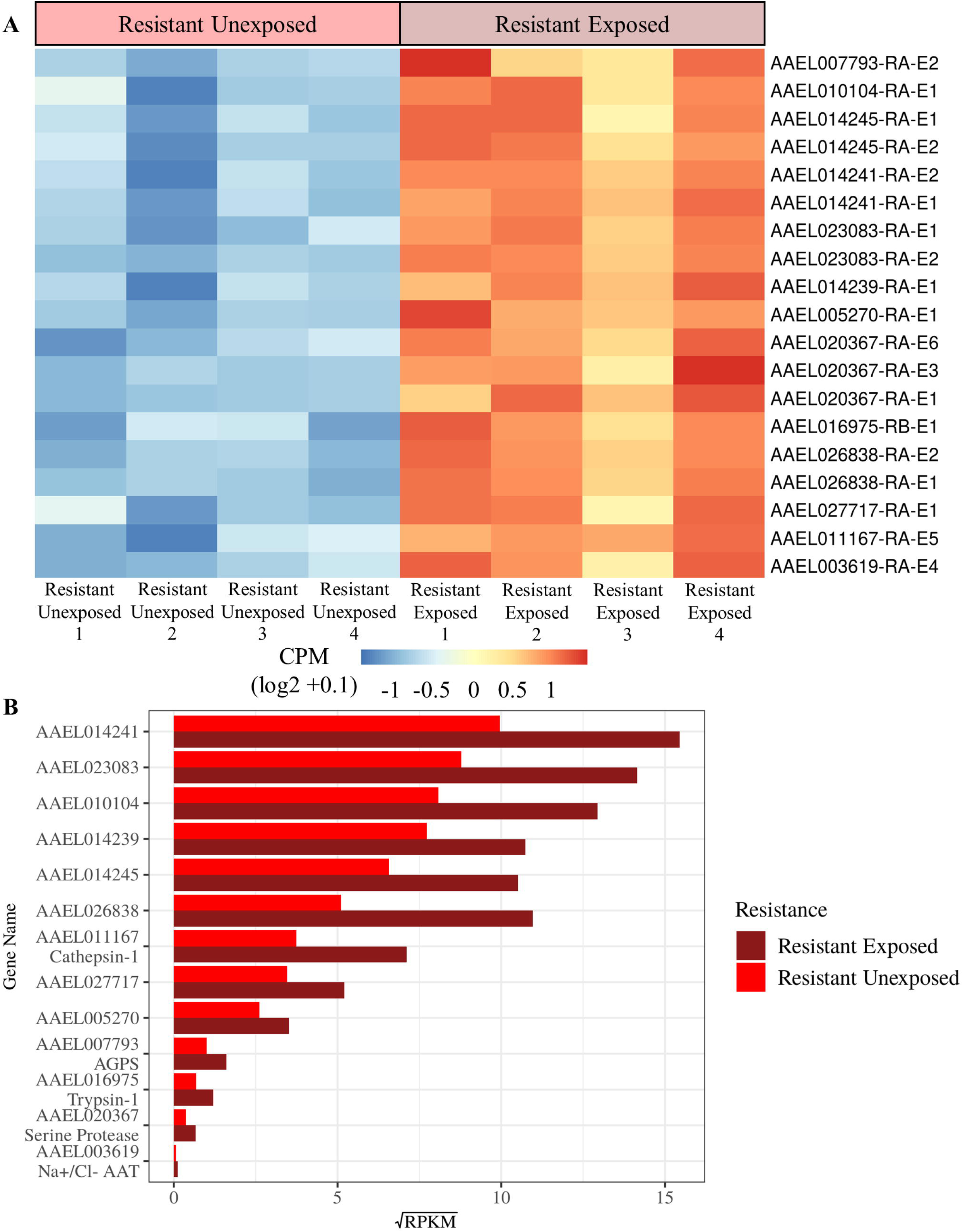
Differential gene expression of temephos-resistant larvae under transient exposure to temephos. Comparison of the resistant temephos exposed population with the resistant unexposed population identified 19 significantly over expressed transcripts and 13 significantly over expressed genes. The transcript expression was displayed here (A) as counts per million (CPM) which is the number of counts per transcript following normalisation. Expression was displayed per sample, rather than per experimental group, allowing for visualisation of granularity between samples. The expression was scaled by row to allow comparison between samples rather than between transcripts. Variability in expression between samples from the same experimental groups can be seen across all genes, highlighting the importance of biological replication. Gene expression was displayed here (B) as reads per kilobase million (RPKM) in the resistant exposed population (darker red) and resistant unexposed population (lighter red). RPKM values were calculated for each gene by taking the mean RPKM of each transcript within that gene. The RPKM values were square root transformed here to optimise the visualisation of a vast range of values (0.003 to 239). Mean RPKM values per group allow for comparison of expression between genes as well as between groups.

## Discussion

Management of arbovirus burden is threatened by insecticide resistance in mosquitoes which reduces the effectivity of vector control [70–72]. In this study we report resistance to temephos in the field population of *Ae. aegypti* from Cúcuta whilst larvae from Bello were susceptible. Bello is an area of relatively low arbovirus incidence[44] and has a lower frequency of insecticide usage [73], whilst Cúcuta is an area of high arbovirus incidence which has seen routine use of temephos for *Ae. aegypti* control over four decades [17]. The reported resistance in Cúcuta is consistent with previous reports of temephos resistance in *Ae. aegypti* from Cúcuta in 2010, seven years earlier than the mosquito collections took place for this current study [17]. Whilst the resistance to temephos appears to have reduced in Cúcuta from RR_50_ = 11.85 in 2010 [17] to RR_50_ = 8.0 in 2017 (current study) resistance to temephos remains moderate demonstrating the long-term implications of insecticide resistance on vector control programs. Management of arbovirus burden is threatened by insecticide resistance in mosquitoes which reduces the effectivity of vector control programs [71, 74], including alternatives such as biological control strategies [72].

The triangulation of differential gene expression against two unrelated susceptible populations, one lab and one field, was selected to reduce confounding effects of phenotypic differences between populations unrelated to insecticide resistance. Whilst this experimental design does reduce these confounding effects it is not possible to mitigate this entirely and therefore some of the differences in gene expression which are observed here may not be related to temephos resistance but resistance to other insecticides and other phenotypic differences between populations. The differential gene expression reported here could be the reflection of the selective pressure under other larvicidal insecticides used in Cúcuta in a similar time span and even selective pressure from adulticides, such as malathion, fenitrothion, λ-cyhalothrin and deltamethrin [6, 16], through vertical transfer. *Ae. Aegypti* larvae from Cúcuta have previously been reported to be highly resistant to the pyrethroid -λ cyhalothrin whilst larvae from Bello were susceptible [75]. These findings from the same study localities used in the current study demonstrate the effect adulticide resistance can have on larvae. Cross resistance between organophosphates, such as temephos and pyrethroids has also been reported in *Ae. aegypti* [76–78], including in Colombian *Ae. aegypti* populations [16].

Differential gene expression associated with the resistant phenotype was identified by selecting genes which were differentially expressed in the field resistant population compared to both the field (Bello) and lab (New Orleans) susceptible populations. This reduced the cofounding effects of location differences and enabled the analysis to focus on DGE associated specifically with resistance. This comparison identified 503 significantly differentially expressed genes which are potentially associated with the resistant phenotype, 301 of which were over expressed in the resistant population and 239 under expressed.

Genes which were found to be differentially expressed in the current study may also be the result of epistatic interactions, genetic and biochemical, and therefore associated with other biological processes aside from insecticide resistance, such as those which compensate for resistance induced fitness cost [79–81]. Epistatic interactions between genes associated with insecticide resistance are also known to influence different levels of resistance [82]. Under such conceptual framework the following functional categories are highlighted.

### Metabolic detoxification genes

Metabolic detoxification of insecticides is one of the most reported insecticide resistance mechanisms in mosquitoes. Abundant overexpression of detoxification genes, most commonly cytochrome P450 monooxygenases (P450), glutathione S-transferases (GST) and carboxylesterases (CE) has frequently been associated with insecticide resistance in mosquitoes [83]. Here we reported the overexpression of only two P450s (*CYP12F6 -* AAEL002005 and *CYP6BY1 -*AAEL017539), one GST (AAEL006818) and one CE (*CCEAE4C* (AAEL003187)), in the resistant population compared to both field and lab susceptible populations. CYP12F6 has previously been shown to be overexpressed in a permethrin resistant population of adult *Ae. aegypti* from Mexico albeit compared with a lab susceptible population only [41]. GO terms associated with insecticide detoxification (oxidoreductase activity (GO:0016491) and oxidation-reduction process (GO:0055114)) were also found to be enriched in the temephos resistant larvae. Thus, by cross examining the data in field-to-field and field-to-lab population comparison, we observed genes representing these three forms of insecticide deactivation in much reduced number compared to what is commonly reported [17,30,59,60]. To illustrate the above, if the resistant population had been compared with the lab susceptible population only a total of 49 cytochrome P450s, six GTSs and 11 CEs would have been reported as differentially expressed (S3 Table). This suggests that large overexpression of detoxification genes may be partly related to differences between field and lab mosquitoes rather than associated with the insecticide resistant phenotype. Large overexpression of detoxification in mosquitoes may also only be observed in mosquitoes when they have high levels of resistance rather than the moderate resistance reported here [59, 84].

### Chitin biosynthesis

The thickness and composition of the cuticle has been identified as a critical determinant of insecticide resistance due to its role in reducing insecticide penetration [33]. Over expression of genes associated with formation and maintenance of the cuticle have been reported in insecticide resistant populations of medically relevant species including *An. gambiae* [85–87], *An. funestus* [88] and *Culex pipiens pallens* [89, 90]. The cuticle has also been associated with resistance in *Ae. aegypti* including in larvae [34, 36]. The over expression of the chitin biosynthesis enzyme AAEL002718 and the enrichment of chitin synthase activity (GO:0004100) in temephos resistant *Ae. aegypti* larvae reported in this study further highlights the potential role of the cuticle in the development of insecticide resistance in *Ae. aegypti* larvae. Chitin, a biopolymer of N-acetylglucosamine, is a major constituent of the mosquito cuticle (exoskeleton (epidermal cuticle), tracheal cuticle and eggshell) providing it with both strength and rigidity and is also found in midgut peritrophic matrices [91]. The use of chitin synthesis inhibitors (CSI), a type of insect growth regulators (IGRs) which interfere with the synthesis and deposition of chitin on the exoskeleton [92], has been highlighted as a potential approach to control *Ae. aegypti* with some promising findings in laboratory studies [93, 94].

### Pattern recognition and innate immunity

Temephos resistant *Ae. aegypti* larvae were shown to express high levels of the mannose-binding C-type lectins (CTLs) AAEL011612 and AAEL000533 which are predominantly produced in the salivary glands of adult female *Ae. aegypti* [95, 96]. Lectins are ubiquitous proteins in multicellular organisms that provide the pattern recognition required for the initial phase of an immune response [67, 68]. C-type lectins are a group of calcium-dependant carbohydrate binding proteins [97]. In mosquitoes CTLs are primarily involved in facilitating viral infection (e.g., dengue, Rift Valley fever and Japanese encephalitis viruses [98, 99]) through enhanced viral entry, acting as bridges between flaviviruses and host cell receptors [99, 100]. However, these proficient pattern recognition proteins seem to have evolved to mediate multiple multicellular processes beyond mosquito immune response including lifespan and reproductive capability [101] as well as maintenance of gut microbiome homeostasis [102]. Transferrin (*Tf1* - AAEL015458), found to be under expressed in temephos resistant larvae in the current study, also has roles in the innate immune response to arbovirus infection [98, 103] and has previously been reported to be downregulated in CHIKV and DENV infected mosquitoes which may favour viral replication [104]. Transferrin expression has also been related to insecticide resistance in *Culex pipiens* with increased expression reported in mosquitoes with target-site resistance to pyrethroids and organophosphates, the biggest difference in transferrin expression was observed in adults [105, 106].

### Cell membrane transport

The expression of several ion coupled solute membrane transporters was down regulated in temephos resistant larvae: the sodium-coupled cation-chloride cotransporter AAEL009886 (aeCC3), the sodium/chloride dependent amino acid transporter AAEL000298, the sodium/solute symporter AAEL001198, and the sugar transporter AAEL010348. aeCCC3 is a larvae specific membrane transporter abundant in the anal papillae responsible for the absorption of external ions [107] which belongs to a family of cation-coupled chloride cotransporters (CCCs) which contribute to ion homeostasis by undertaking electroneutral transport of Na^+^, K+ and Cl^-^ [108]. A similar role is expected from the ion-coupled transporters AAEL000298 and AAEL001198 in the homeostasis of ion content, particularly in midgut and Malpighian tubes where they are most abundant [109].

The aquatic life of the *Ae. aegypti* larval stages demands an ion exchange homeostasis that differs from that of the adult mosquitoes. Due to their freshwater habitat *Ae. aegypti* larvae must excrete water gained by osmosis, reabsorb salt prior to excreting urine, and absorb salt from their surroundings [110]. Whilst the opposite is true in adults where water retention is needed due to constant loss through evaporation. A key process in this is Na+-dependent co-transport which is typically down the large inward (extracellular to intracellular) Na+ gradient generated by the Na+/K+-ATPase [111]. We speculate that the ion homeostasis changes caused by the reduced expression of the three CCCs transporters AAEL009886, AAEL000298 and AAEL001198 could reduce the exposure of larvae to temephos by reducing net uptake of the molecule, protecting the organs where they are commonly expressed (e.g., midgut and Malpighian tubes). Transcriptome studies of insecticide resistant mosquito populations tend to overlook the potential role of down regulated genes in favour of overexpressed genes, but this finding demonstrates the importance of investigating reduced expression when studying potential mechanisms of insecticide resistance. The potential role of CCC transporters in reducing insecticide uptake and therefore facilitating resistance warrants further investigation.

### Chromosomal remodelling

The mdg4-binding protein ortholog gene (AAEL010576: modifier of mdg4 [Mod(mdg4)]), responsible for chromosome remodelling was also significantly under expressed in the resistant *Ae. aegypti* larvae. Originally described as a protein binding the transposon mdg4 [112], Mod(mdg4) gene encodes for a family of proteins due to at least 30 different alternative splicing variants in Diptera and Lepidoptera [113–115]. Mod(mdg4) variants bind a variety of insulators (DNA domains involved in nuclear organization and higher order chromatin structures) [116–118] and have been involved in regulating numerous traits of the insect embryonic progression such as synapsis structure [119], chromosome Y-linked testis development [120], and mid-gut maturation [121]. Changes in expression of Mod(mdg4) have been reported in *Drosophila* Kc cells treated with deltamethrin [122]. The identification of under expression of the mdg4-binding protein in temephos resistant larvae suggests a further role for this protein in mediating insecticide resistance.

### Long non-coding RNA

There were 55 over expressed and 30 under expressed genes encoding for long non-coding RNA (lncRNA) in the temephos resistant larvae (S1 Table). Non-coding RNAs (ncRNA) are abundant cellular effectors of great prolific functionality [123] and long ncRNA are defined as transcripts, more than 200 nucleotides long, that are produced by RNA polymerase II and are not translated into proteins [124]. In *Aedes* lncRNAs, mainly involved in regulating gene expression, are multifunction with roles including sex differentiation [125], embryogenesis [126] and suppression of viral replication in DENV infected mosquitoes [127]. Long ncRNAs have also been associated with insect’s response to xenobiotics, with reports of differential lncRNA expression in resistant populations of *Plutella xylostella* [128]. The findings of 85 differentially expressed lncRNAs reported here in resistant populations of *Ae. aegypti* supports the potential roles that lncRNAs could have in the development of insecticide resistance. Whilst gene expression studies have focussed primarily on differential expression in protein coding genes, the development of next generation techniques have now provided an opportunity to also study noncoding RNA. Whilst work has been conducted into identifying lncRNAs in medically relevant mosquito species including *An. gambaie* [129] and *Ae. aegypti* [126] there have been no studies that have aimed to investigate the role of lncRNAs in insecticide resistance in mosquitoes. Previous RNA-Seq studies on insecticide resistant populations of *Culex pipiens pallens* have also identified differential expression of lncRNAs [130], however, an in-depth discussion of their role in insecticide resistance has been neglected.

### Differential gene expression in resistant larvae following temephos exposure

In the study we also tracked gene expression in insecticide resistant larvae following direct response to temephos exposure. Thirteen genes were found to have a significantly increased expression following a controlled exposure to temephos. Among those 13 genes were two serine proteases: trypsin -1 (AAEL016975) and serine protease stubble (AAEL020367), a cysteine protease: cathepsin-1 (AAEL011167), a sodium/chloride dependent amino acid transporter (AAEL003619) and an alkyl dihydroxyacetone phosphate synthase (AAEL007793). Serine proteases are a group of enzymes with a variety of known functions including digestion, metamorphosis, oogenesis, blood coagulation and viral immune response [98,131,132]. Cathepsin-1 (AAEL011167), a cysteine proteinase, is also a multifunctional digestive enzyme [131, 133]. Upregulation of serine proteases have been previously reported in insecticide resistant mosquito populations [39,41,134,135], including temephos resistant *Ae. aegypti* from Cúcuta [17]. Serine proteases have also been shown to degrade insecticides through hydrolysis within the insect digestive tract, however so far evidence of this is limited to pyrethroids such as deltamethrin [136–139]. Overexpressed proteases in this current study support the findings of previous studies that proteases may have a role in the metabolism of other insecticide classes besides pyrethroids. The changes responsible for resistance are often associated with modification of physiological processes that can lead to decreased performance and fitness disadvantage.

Deleterious effects of insecticide resistance can affect a wide range of life-history traits (e.g. longevity, biting behaviour, and vector competence) [140, 141]. Although the cost of resistance genes is believed to gradually decrease due to subsequent modifier mutations [142]. With the relatively limited diversity of insecticide targets [143], the gene expression patterns that resistant mosquitoes further undergo when exposed to the insecticide could be a source for novel assets for vector control. The study of such targets for insecticide development is a strategy that, to our knowledge, has not yet been explored.

## Conclusion

This study found differential insecticide responses from *Ae. aegypti* field samples of two previously epidemiologically characterised sites in Colombia. Using these contrasting *Ae. aegypti* field mosquito populations together with the New Orleans lab strain, we demonstrated the risk of producing noise signal by overestimating by several fold the differential gene expression if mosquito populations are compared only with laboratory strains. The two overexpressed P450s in resistant *Ae. aegypti* larvae represent some ten-fold lower levels in comparison to previous studies [59, 84]. The role of the cuticle in insecticide resistance suggested in previous studies is substantiated here. This study identified other potential mechanisms not previously associated with insecticide resistance in mosquitoes. These included changes in ion exchange homeostasis, chromatin remodelling, lectin-mediated immune responses, and a plethora of lncRNAs. Evidently, there is a notorious gap in our knowledge base of gene expression adaption in insecticide resistance. The work presented here contributed to what seems to be an expansive and varied phenotypic landscape in the *Ae. aegypti* responses to insecticides of current importance.

## Acknowledgements

The work was funded by British Council Institutional Links Newton Fund and supported by the project Research Infrastructures for the control of vector-borne diseases (Infravec2), which has received funding from the European Union’s Horizon 2020 research and innovation program under grant agreement No 731060. The authors thank The University of Liverpool for the support to J.E.S-S, Edge Hill University for support to JM and CS and Polo d’Innovazione Genomica, Genetica e Biologia (poloGGB) for conducting the sequencing.

## Supporting Information Captions

**S1 Table: Significantly differentially expressed genes and transcripts in FR samples when compared to FS and LS samples.** A total of 379 transcripts covering 301 genes were significantly over expressed and 244 transcripts covering 202 genes were significantly under expressed in the resistant population when compared to both susceptible populations. Genomic location, product description, gene type and gene name/symbol obtained from VectorBase annotations. Reads per kilobase million (RPKM) for each population, fold change (logFC), counts per million (logCPM), F-test statistic (F), p value and false discovery rate (FDR) calculated using edgeR.

**S2 Table: Significantly differentially expressed genes and transcripts in the resistant population following temephos exposure.** A total of 19 transcripts covering 13 genes were significantly over expressed in the temephos exposed resistant population when compared to unexposed resistant larvae. Genomic location, product description, gene type and gene name/symbol obtained from VectorBase annotations. Reads per kilobase million (RPKM) for each population, fold change (logFC), counts per million (logCPM), F-test statistic (F), p value and false discovery rate (FDR) calculated using edgeR. Uncharacterised genes were searched for homologs in other species using NCBI nucleotide blast (https://blast.ncbi.nlm.nih.gov/Blast.cgi) but no characterised homologs were identified.

**S3 Table: Significantly differentially expressed genes and transcripts in FR samples when compared to FS samples only.** A total of 3328 transcripts covering 2322 genes were significantly over expressed and 2250 transcripts covering 1555 genes were significantly under expressed in the resistant population when compared to the lab susceptible population. Genomic location, product description, gene type and gene name/symbol obtained from VectorBase annotations. Reads per kilobase million (RPKM) for each population, fold change (logFC), counts per million (logCPM), F-test statistic (F), p value and false discovery rate (FDR) calculated using edgeR.

